# genepanel.iobio - an easy to use web tool for generating disease- and phenotype-associated gene lists

**DOI:** 10.1101/722843

**Authors:** Aditya Ekawade, Matt Velinder, Alistair Ward, Tonya DiSera, Gabor Marth

## Abstract

A comprehensive list of candidate genes that succinctly describe the complete and objective phenotypic features of disease is critical when both ordering genetic testing and when triaging candidate variants in exome and genome sequencing studies. Great efforts have been made to curate gene:disease associations both in academic research and commercial gene testing settings. However, many of these valuable resources exist as islands and must be used independently, generating static, single-resource gene:disease association lists. To more effectively utilize these resources we created *genepanel.iobio* (https://genepanel.iobio.io), an easy to use, free and open-source web tool for generating disease- and phenotype-associated gene lists from multiple gene:disease association resources, including the NCBI Genetic Testing Registry (GTR), Phenolyzer, and the Human Phenotype Ontology (HPO). We demonstrate the utility of *genepanel.iobio* by applying it to complex, rare and undiagnosed disease cases which had reached a diagnostic conclusion. We find that *genepanel.iobio* is able to correctly prioritize the diagnostic variant in over half of these challenging cases. Importantly, each resource contributed diagnostic value, showing the benefits of this aggregate approach. We expect *genepanel.iobio* will improve the ease and diagnostic value of generating gene:disease association lists for genetic test ordering and whole genome or exome sequencing variant prioritization.

## Background

A tremendous amount of genetic and biomedical knowledge has been deposited in both curated and computationally-derived gene:disease association database resources such as NCBI’s Genetic Testing Registry (GTR) (Rubinstein et al. 2013), Phenolyzer (Yang, Robinson, and Wang 2015), and the Human Phenotype Ontology (HPO) (Robinson et al. 2008). However, each resource has its own format, data structure and output, making it difficult for a genetics professional to navigate between resources and especially difficult to merge the outputs of these resources into a concise, non-redundant list that encompasses the phenotypic features of disease. As such, there is a strong need in the medical genetics community for a tool capable of consolidating and harmonizing these gene:disease association resources and providing genetics professionals with a single prioritized candidate gene list.

## Implementation

Our primary goal in developing *genepanel.iobio* was to make it as universally accessible as possible, ensuring clinicians and genetics professionals without prior computational expertise could adopt it into their clinical workflows. This goal lent itself towards a web-browser application capable of being run on any computer, regardless of operating system or hardware specifications and reflects our goals for the *iobio* platform more generally (Ward et al. 2017). Considering this, we built *genepanel.iobio* to fully utilize the modern progressive component-based Javascript framework, Vue. Vue allows us to write, reuse and share components for creating a platform-independent application that runs entirely in a web browser. Within this framework we can fetch data from various application programming interfaces (APIs) such as the NCBI’s Genetic Testing Registry (Figure 1A). Querying data via an API ensures the *genepanel.iobio* user is retrieving the most up-to-date information available. For tools without an available API (Phenolyzer and ClinPhen) we developed backend services, written in Node.js, that create a wrapper around the original command-line tools. We optimized these backend services, for example, by caching Phenolyzer’s output in DynamoDB (a NoSQL database), allowing for near instantaneous results of previously searched terms. The application is hosted on AWS S3 and utilizes AWS Cloudfront services. This provides a fast content delivery network (CDN) which is massively scaled and globally distributed with superior performance and stability. The application is also configured with HTTPS to ensure the user has secure end-to-end connections to the origin servers. Numerous other design considerations were implemented based on clinician feedback, including: implementing typeahead prediction for terms; changes to the user-interface; implementing advanced filtering for each resource; adding additional resources such as HPO; implementing export features; and displaying pertinent gene information for each gene. On the whole, this approach has allowed us to harmonize disparate data sources, structures and programs into a single easy-to-use, broadly adoptable application. Our code is free, open-source and available at https://github.com/iobio/genepanel.iobio.

**Figure 1:**
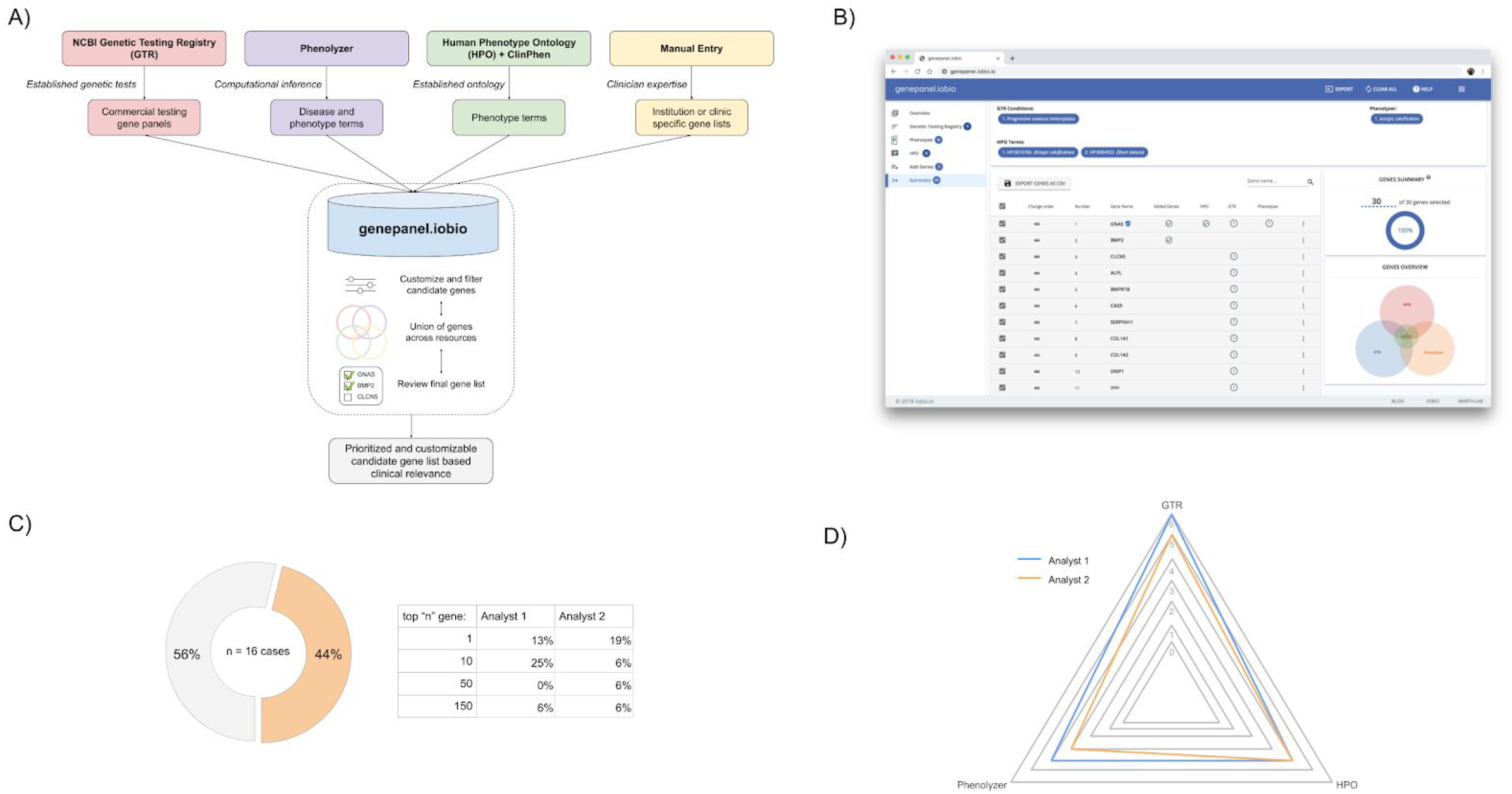
An overview of *genepanel.iobio* applied to clinical genetics cases A) a diagrammatic view of the resources *genepanel.iobio* utilizes and how *genepanel.iobio* prioritizes a unioned set of candidate genes B) a screenshot view of the summary page of a *genepanel.iobio* analysis, showing the ability to modify and customize the final gene list and export the results C) two independent analysts correctly prioritize the diagnostic gene using *genepanel.iobio* in 7 of the 16 clinical cases (44%) tested and where the diagnostic gene was ranked in the final gene list by each analyst D) radar-plot summary of which resources correctly prioritized the diagnostic gene, allowing for multiple resources to identify the same gene

## Results/Discussion

Within a typical *genepanel.iobio* usage, a user provides relevant terms to each resource tool. For instance, starting with the NCBI Genetic Testing Registry input step, the user will enter one or more presumed genetic disorder terms, selecting the desired term from the typeahead drop down. Proceeding through the Phenolyzer input step, the user will search and select one or more disease-relevant phenotype terms. In the HPO input step the user can either directly input HPO terms or copy/paste a clinical note and allow the ClinPhen (Deisseroth et al. 2018) computational tool to identify and rank relevant HPO terms from the clinical note (Figure 1A). These input steps can be performed independently of one another, allowing the user to utilize some or all of the available resources. Within each input step the user can apply advanced resource-specific filters to further customize and refine the genes generated by that resource. For example, in the GTR step the user can filter on commercial testing providers and/or disease modes of inheritance. Following data input *genepanel.iobio* presents the user with a summary page of the prioritized union of genes across all resources. This final step allows the user to further refine the gene list and come to consensus as to the most disease relevant genes (Figure 1B). The user can then export the final gene list into a text file, comma-separated file or copy them to the system clipboard.

Following development, we tested the efficacy of *genepanel.iobio* to correctly prioritize diagnostic variants in a clinical genetics setting. We chose an ambitious test setting, selecting cases from a rare and undiagnosed disease genetics clinic at the University of Utah which had previously reached a diagnostic conclusion. The rate of diagnosis in this setting remains between 35-45%, often complicated by complex clinical presentations and ambiguous or blended phenotypes (Demos et al. 2019; Deignan et al. 2019). Two analysts, blinded to the causal variants, independently analyzed 16 previously diagnosed clinical cases to see if *genepanel.iobio* could correctly prioritize the diagnostic gene within the final gene list. These analysts had no speciality or expert knowledge of the genetic disorders or the genes associated with them. These analysts were able to correctly prioritize the diagnostic gene in their final gene list in 7 of the 16 clinical cases (44%). The total number of genes in the final gene list were typically less than 150 genes, a number we determined to be reasonable for gene panel testing and genetic sequencing variant review workflows. The diagnostic gene was the number one ranked gene in the final gene list for 2 (analyst 1) and 3 (analyst 2) of the clinical cases. Additionally, for over one third of cases (44% analyst 1, 38% analyst 2), the diagnostic gene was in the top 50 genes of the final gene list (Figure 1C). Interestingly, each resource contributed to the analysts’ ability to correctly prioritize the diagnostic gene (Figure 1D). Additional metrics about the analysts gene list generation can be found in Supplemental Table 1.

These results demonstrate the benefit of *genepanel.iobio* and its aggregate approach of using multiple gene:disease association resources. We are actively developing and maintaining *genepanel.iobio* and will be incorporating new features and resources in the future. We predict that additional resources will only further add to the power of *genepanel.iobio*.

## Conclusion

Genomic medicine has greatly benefited from the increasing wealth of knowledge in gene:disease association databases and resources. However, it remains difficult to harmonize results across multiple gene:disease association resources. To address this difficulty we developed *genepanel.iobio*, a free, open-source, platform independent web application capable of generating a comprehensive list of genes associated with a user-provided set of phenotypes. We demonstrate the utility of *genepanel.iobio* in a clinical genetics setting by its ability to generate a gene list containing a reasonable number of genes that encompass the clinical phenotype and mostly importantly contained the diagnostic gene. We anticipate adoption of *genepanel.iobio* into clinical genetics workflows would improve the diagnostic value of genetic test ordering as well as variant/gene prioritization in genetic sequencing studies.

## Supporting information

figure1_analyst_details

